# Nanotopographic Control of Actin Waves and Growth Cone Navigation in Developing Neurons

**DOI:** 10.1101/2025.05.16.654525

**Authors:** Spandan Pathak, Kate M. O’Neill, Emily K. Robinson, Matt J. Hourwitz, Corey Herr, John T. Fourkas, Edward Giniger, Wolfgang Losert

## Abstract

The development of axons and dendrites (neurites) in a neural circuit relies on the dynamic interplay of cytoskeletal components, especially actin, and the integration of diverse environmental cues. Building on prior findings that actin dynamics can serve as a primary sensor of physical guidance cues, this work investigates the role of nanotopography in modulating and guiding actin waves and neurite-tip dynamics during early neural circuit development. Although actin dynamics is well known to contribute to pathfinding in wide axonal tips, typically referred to as growth cones, we also observe dynamic actin remodeling throughout neurites and at other, narrower, neurite tips. We find that actin-wave speeds do not change significantly in the first two weeks of neurite development on flat substrates, but decrease over the same period in neurites on nanoridges. The ability of nanoridges to guide actin waves and the neurite-tip direction also decreases as neurites mature, both for narrow tips and wide growth cones. This change in responsiveness to physical guidance cues with neuronal maturation may impact the regenerative capacity of developing neural cells that are inserted into mature brains.

## 1. Introduction

The development of a functional nervous system is a marvel of Nature, relying on the precise navigation and connection of countless neurons. This intricate process of neural circuit formation relies on the dynamic behavior of developing neurons that extend neurites – projections that will ultimately become axons and dendrites (1, 2). This navigation depends critically on the active and tightly regulated remodeling of the actin cytoskeleton (3, 4). Actin filaments (F-actin), in conjunction with microtubules (5, 6), underlie the extension and steering of axons, as well as the formation of synapses (7).

Actin waves in migrating eukaryotic cells have been studied and characterized extensively. Actin waves are domains of locally high actin density that translocate within the cell. They can, for example, be foci that move within a neurite (8, 9), or broad fronts within a cell body (10, 11), and they are wave-like in that their motion reflects translocation of the density distribution and not of the actin monomers themselves. The motion of actin waves arises from the interaction of actin with its associated network of cytoskeletal regulators that forms a system of biomechanochemical excitability and it is deeply involved in cell migration. In the slime mold Dictyostelium discoideum, actin waves have been shown to be responsible for migration (12) and for texture sensing (13, 14). Other members of the signal-transduction network, particularly the actin-related protein 2/3 (Arp2/3) complex and PIP3, also form complex waves with specific localization patterns within a cell (12–15). Interestingly, actin dynamics tends not to be required for signal-transduction waves to occur (16–19), yet both types of waves are necessary for important functions such as cell motility, and demonstrate an intricate interplay of signaling (20).

In developing neurons, actin waves are most commonly observed in association with neurite extension, where these waves precede and promote axon outgrowth. Although growth-cone-led axonal development has been studied widely, recent work has shown that cytoskeletal dynamics may also direct growth for narrow neurite tips (21, 22). It was demonstrated recently, for example, that in the *Drosophila* wing disc, the tip of the pioneer neuron TSM1 is almost exclusively narrow and filopodial (23). In this case, genes for proteins such as Abelson tyrosine kinase regulate actin organization and, therefore, the advance of the axonal tip (23, 24) through actin dynamics that occurs distant from the leading edge. Neurons also seem to possess mechanisms to regulate neurite elongation that act in regions other than the growth cones. For example, differential neurite growth, including the selection of a single axon during initial polarization and the alternating growth of axon branches, occurs even in the absence of growth-cone motility (25). This observation suggests that neurons possess mechanisms for local regulation of neurite elongation in regions other than the growth cones.

The morphology of neurons changes dramatically during development (26). Dendrites begin as small protrusions from the cell body, and grow into a complex network of branches that ultimately determines how a neuron receives input (27). Neurons in an earlier stage of development – which we define as day *in vitro* (DIV) 2 through 5 in our study – are characterized by a simpler morphology, with primary and secondary dendrites and likely to have a specified axon. Neurons in the middle developmental stage, DIV 7 through 12 in our study, have more complex dendritic trees with tertiary dendrites and longer axons. Finally, neurons in a later developmental stage, DIV 14 through 23 in our study, have intricate dendritic structures, and are likely to be developing synaptic spines. Given how neurites (axons and dendrites) change significantly during development, one goal of our study is to ask whether the dynamics of the cytoskeleton also changes during development.

Further, although the actin waves that traverse the length of the axon are associated with growth and retraction, conflicting views have emerged regarding the precise roles of these waves. On the one hand, neurite elongation, which may reach lengths of hundreds of microns, can occur independently of actin waves (28). Conversely, studies have shown that, during early neuronal development, actin waves drive fluctuations in neurite outgrowth by controlling microtubule transport (8). In some cases, competition among neurites, which is mediated by actin waves, is proposed to allow the neuron to explore its environment until external cues specify the axon (29). These findings suggest that actin waves act more as passive responders to stochastic internal forces rather than as active drivers of axon extension and growth-cone formation.

Classic studies of axon patterning, such as those described above, have largely focused on what happens inside the axon and growth cone. However, the physical properties of the microenvironment, such as substrate stiffness (30) and topography, also can exert a profound influence on neuronal growth behavior in different stages of development (31, 32). In early stages (4 to 20 hours post plating), specific topographic cues can direct neuronal polarization by attracting key markers, resulting in preferential axon growth along micropatterned features (33). With further development (4 to 5 days post plating), growth cones on such substrates became more elongated and aligned with respect to the neurite shaft (34). Moreover, substrate stiffness is not only relevant early in development, but also later in the mature brain (35). Studies indicate that region-specific changes to stiffness occur with age, and particularly that the aging brain decreases in stiffness (36, 37). In neurodegenerative diseases, such as Alzheimer’s (38, 39) or Lewy-body dementia (40), brain stiffness is decreased even further in areas relevant to those diseases. These findings highlight the importance of considering the three-dimensional complexity of the *in vivo* environment when studying the role of actin dynamics in axonal guidance, growth-cone navigation, and longer-term neural circuit function.

The current work investigates the role of nanoridges in modulating actin waves in neurites and growth cones during neuronal development. The guidance of neurites by nanoridges has been examined previously (41), and our goal is to characterize the behavior of actin waves under this well-studied mechanical cue (42) and under control conditions. Although nanoridges are a simplified model for variable physical cues, the ridges may mimic pre-existing nerve structures along which follower axons grow, and that are mechanically distinct from the surrounding epithelium. Prior work suggests that axon-guidance mechanisms differ between follower and pioneer axons (43), lending physiological relevance to our experimental setting. The nanoridges we used, had a 3.2 μm spacing, allowing us to compare with our previous work on other cell types (44–48). We hypothesized that this spacing would enable the guidance of neurites (49, 50). Although the nanoridges have a stiffness comparable to that of physiological substrates such as collagen fibers, the ridges are rigidly fixed to the underlying surface, and do not deform. This distinction highlights the fact that nanotopographical cues mimic some, but not all, of the mechanical aspects of the *in vivo* environment. We examine the speed and the guidance of actin waves, neurites, and growth cones by this nanotopography. Because actin waves have been identified as primary sensors of the physical microenvironment, including electric fields (51) and texture (46, 52), our work also sheds light on whether neuronal development also alters the ability of cells to sense their physical environment.

## 2. Materials and Methods

### 2.1 Primary neuronal dissections and cell culture

Primary neurons were obtained from Sprague–Dawley rats housed at the University of Maryland (with approval by the University of Maryland Institutional Animal Care and Use Committee; protocols R-JAN-18-05 and R-FEB-21-04). At embryonic day of gestation 18 (E18), primary cortical neurons were obtained via dissection, and were cultured as previously described (53). We did not distinguish between cortices from male and female embryos. Briefly, hippocampi and cortices were dissected from decapitated embryos. Cortices from all embryos were combined and triturated manually, and then were counted using a hemocytometer. A monolayer of cortical neurons was plated in 35-mm-diameter imaging dishes with glass centers (P35G-1.5-20-C, MATTEK) at a density of 5 × 10^5^ cells per dish (520 cells/mm^2^). Prior to the plating, the imaging dishes were coated with 0.1 mg/mL poly-D-lysine (PDL; P0899, MilliporeSigma) for at least 1 h at 37 °C and then were washed three times with phosphate buffered saline (PBS; SH30256.01, Cytiva). Monolayers were also plated on acrylic nanoridges, which received the following treatments prior to neuronal plating: soaking in ethanol (24 h), followed by soaking in medium (24 h), followed by PDL coating (24 h), and finally 10 μg/ml laminin coating (24 h; L2020, MilliporeSigma), all at 37 °C. Cultures were maintained in NbActiv4 medium (Nb4-500, BrainBits) supplemented with 1% penicillin/streptomycin (P/S; 15070063, Thermo Fisher Scientific) at 37 °C in 5% CO_2_. Half of the medium was changed twice per week. Note that we did not optimize the nanoridges for longer term neuronal survival (DIV14+), which is why our data for cells on nanoridges are only reported at the early and middle developmental stages.

### 2.2 Transduction with Actin-GFP

For visualization of actin dynamics within primary cortical neurons, we used CellLight Actin-GFP, BacMam 2.0 (C10506, ThermoFisher Scientific). This viral vector has low toxicity and labels cells sparsely. Given that neurons have complex morphological structures even in early stages of development, the latter feature is particularly helpful for analysis of single-cell actin dynamics. At 48 to 72 h prior to imaging, 5 to 10 μL of CellLight was mixed with the cell-culture medium (1 to 2 particles per cell). The medium was fully changed approximately 20 h after transduction, and imaging was performed 24 to 48 h later.

We chose to use actin-GFP transduction via a baculovirus. The low toxicity of the baculovirus promotes healthier primary cultures, and the low efficiency of transduction of this vector ensured that we could analyze actin dynamics of individual neurons. However, using transduction of actin-GFP rather than a fluorescent actin dye introduced certain limitations to our study. In particular, our nanotopopgrahic surfaces, although optimized for primary cell survival, are not ideal for longer-term (DIV 10+) neuronal survival. Our preliminary studies of transduced, late stage neurons on surfaces did not yield healthy cultures. Therefore, we chose to limit our studies of neurons on ridges to the early and middle developmental stages, to ensure the integrity of, and consistency within, our results. The combined challenge of low transduction efficiency and primary culture health on non-standard surfaces is also the reason that the number of replicates for experiments on nanotopographic surfaces is lower than on control surfaces.

### 2.3 Confocal imaging

Live-cell imaging of primary cortical actin dynamics was performed at the University of Maryland Imaging Core using a PerkinElmer spinning-disk confocal microscope. For all experiments, we used an oil-immersion, 40× objective (1.30 NA; 0.36 μm/pixel), and maintained live cultures under temperature, humidity, and CO_2_ control. We recorded 16-bit images using a Hamamatsu ImagEM X2 EM-CCD camera (C9100-23B) and acquired time-lapse images with PerkinElmer’s Volocity software (v. 6.4.0). We identified transduced neurons via a positive signal in the 488-nm excitation channel. We performed imaging in growth medium and acquired images every 2 s in the 488-nm channel for 5 to 10 min. We ensured a consistent *z*-plane with Perfect Focus (Nikon PFS).

### 2.4 Preparation of nanoridges

Nanoridged substrates for the neuronal experiments featured the same master pattern as in a recent study that focused on the photomodification and reshaping of nanotopographic structures (54). Briefly, a large-scale pattern with 750-nm high, 400-nm wide nanoridges at a 3.2-μm pitch (54), initially fabricated on a silicon wafer using interference lithography, was replicated using soft lithographic techniques, as described in more detail elsewhere (55), to produce a polydimethylsiloxane (PDMS) mold with negative relief of the master pattern. Molding of the patterned silicon wafer was facilitated by exposure to plasma treatment in an oxygen atmosphere and subsequent surface functionalization of the wafer with (tridecafluoro-1,1,2,2-tetrahydrooctyl) methyldichlorosilane (Gelest, CAS #: 73609-36-6) under vacuum inside a desiccator. These steps reduce the surface energy of the wafer to prevent adhesion with the PDMS during the molding process. A hard PDMS layer was spin-coated onto the pattern and Sylgard 184 (Dow) comprised the remainder of the mold, based on previous work. (55–57). The mold was peeled from the master after several baking steps. The hard-PDMS was allowed to cure at room temperature for 2 h before baking at 60 °C for 1 h. After the Sylgard 184 elastomer base and curing agent were mixed (10:1 ratio) and poured atop the hard-PDMS layer, the composite was baked an additional 70 min at 60 °C. At this point, the mold was peeled from the master. Replicas of the original pattern were then produced using nanoimprint lithography. A photocurable resin was prepared using a 1:1 w/w mixture of ethoxylated(6)trimethylolpropane triacrylate (Sartomer, CAS #: 28961-43-5) and tris (2-hydroxy ethyl) isocyanurate triacrylate (Sartomer, CAS #: 40220-08-04), and 3 wt% of the photoinitiator Irgacure TPO-L (BASF, CAS #: 84434-11-7). Glass coverslips featuring pendant acrylate groups were produced by plasma treatment under oxygen for 3 min followed by stirring in a 2% solution of (3-acryloxypropyl) trimethoxysilane (Gelest, CAS #: 4369-14-6) for at least 12 h. A drop of the resin was sandwiched between the PDMS mold and an acrylated coverslip, and then was exposed to UV light for 5 min to transfer the pattern to the photoresist. The patterned coverslip was then detached from the mold.

### 2.5 Image Analysis

#### Determination of actin-flow vectors using optical flow

In this paper, “actin activity” refers to dynamic actin-polymerization events, which are visible as transient increases in actin fluorescence. These intensity changes propagate in space and time, and are tracked as distinct movement trajectories to quantify single-cell actin dynamics. To prepare the videos for analysis, image-registration and intensity-adjustment functions from the `scikit-image` module (Python) were used to correct for stage drift (jitter) and photobleaching, respectively. To reduce random background noise, processed videos were smoothed temporally using a Gaussian-like filter applied across five successive frames (58). The smoothing timescale was considerably shorter than the timescale of detectable actin-wave propagation, ensuring preservation of relevant dynamics. Subsequently, optical flow was performed on the smoothed image frames using the Lucas-Kanade method (59). A Gaussian weighting matrix (σ = 2 pixels = 0.72 µm) was applied around each pixel location to extract local flow vectors. A reliability threshold of 0.1 was used to remove noisy flow vectors. We verified that varying the size and shape of the Gaussian filter, as well as making modest changes in the reliability threshold, did not significantly affect motion capture or downstream analyses.

#### Neuronal masking

`Scikit-image` (Python) functions were used for the generation and refinement of binary masks. Mean frames were generated, and low-contrast frames were processed using contrast-limited adaptive histogram equalization. A white top-hat filter (with a disk-shaped structuring element of radius = 15 pixels = 5.4 µm) was applied to highlight darker neuronal processes. Adaptive local thresholding (the Niblack method (60)) yielded an initial binary mask, which was refined by multiple iterative morphological opening steps, retaining the *n* largest connected components after manual review and background-noise exclusion. Subsequently, manual selection of a region corresponding to the cell body was employed due to the inherent lack of consistent alignment of the cell bodies. This approach enabled the isolation of neuronal processes from the larger cell mask, and ensured that subsequent analyses focused on the dynamic behavior of actin within the neurites. Neuronal process orientation was determined using a rotating Laplacian-of-Gaussian filter on the mean image-frame (61).

#### Clustering of optical-flow vectors

Optical flow vectors were spatially clustered by quantifying their local alignment with neighboring vectors. For each pixel, we computed the dot product between its flow vector and the average flow vector within an 11 × 11 Gaussian-weighted neighborhood (σ = 5 pixels ≈ 1.8 µm). This enhanced locally coherent motion patterns by reinforcing flow directions that were aligned with their surroundings. (62). Cluster peaks were identified with the `trackpy` (Python) module using an approximate detection diameter of 5 pixels = 1.81 µm and a minimum separation of 15 pixels = 5.4 µm between two neighboring peaks. Detected particle positions for each frame were stored, and were subsequently linked across frames to form trajectories with a search range of 5 pixels = 1.8 µm between successive frames and a memory of one frame (2 s), enabling cluster-track analysis. To emphasize longer-scale actin flows, only tracks persisting for at least eight frames (14 s) were analyzed. We computed and stored both per-frame and per-track metrics. Per-frame metrics included the instantaneous displacement vector, the movement angle, and the instantaneous speed (magnitude of displacement per frame). Per-track scalar metrics included the total track length (the sum of frame-to-frame displacements), the duration (the number of frames), the net displacement (the straight-line distance between the starting and ending points), the average speed (the total track length divided by the duration), the net velocity (the net displacement divided by the duration, reflecting directional persistence), and the sinuosity (the ratio of the total track length to the net displacement). A summary of how these metrics vary with developmental stage and substrate type is presented in Suppl. Fig. 1. In all our downstream analysis, we weighted the histograms of the actin-track metrics by the track duration, so that longer-lasting events contributed more to the distributions. To exclude static clusters, only tracks with an average instantaneous speed greater than 0.5 pixels/frame = 5.4 µm/min were retained.

#### Recurrence

To quantify how consistently new tracks emerged in similar spatial locations over time, we computed the mean spatial recurrence of track start positions across successive time windows. A track’s start position was defined as its earliest detected (*x*,*y*) coordinate. The full time-lapse was divided into non-overlapping windows of a fixed frame duration and within each window, the spatial coordinates of newly appearing tracks were extracted. For each pair of adjacent windows (win1, win2), we identified recurrent tracks as those in win2 whose start positions were within a fixed spatial radius of any start position in win1. The recurrence fraction was computed as the number of recurrent tracks in win2 divided by the total number of new tracks in win2.

To ensure statistical robustness, windows with fewer than a prespecified minimum number of tracks (e.g., 5) were excluded. For each condition, the mean recurrence was calculated by averaging the recurrence fractions across all valid window pairs. The standard error of the mean (SEM) was computed as the standard deviation of these fractions divided by the square root of the number of valid pairs and is reported as error bars in figures.

#### Growth-cone orientation

Growth-cone boundaries were identified manually and were extracted using ImageJ. The alignment of the central shaft was then determined by identifying its major axis.

#### Wasserstein distance

The Wasserstein distance (*W₁)* was used to quantify the dissimilarity between two probability distribution functions (PDFs) (63). This metric represents the area between the corresponding cumulative distribution functions (CDFs). Mathematically, for two probability measures *μ* and *ν* on the real line with CDFs *F* and *G*, respectively, the Wasserstein distance (*W₁)* is defined as *W₁(μ, ν) = ∫ |F(x) - G(x)| dx.* The Wasserstein distance is a useful tool for quantifying modest changes in broad distributions.

#### Pooling of DIV (days *in vitro* after plating) videos and statistical testing

For dynamics analysis, on flat surfaces we categorized videos from DIV 2 to 5 as ‘early’ (*N* = 17), from DIV 7 to 12 as ‘middle’ (*N* = 21), and from DIV 14 to 23 as ‘late’ (*N* = 17) developmental stages. On nanoridges, DIV 2 to 4 were grouped as ‘early-stage’ (*N* = 9), and DIV 7 to 8 as ‘middle-stage’ (*N* = 6).

To compare multiple independent test groups, we first performed the Shapiro-Wilk test to assess normality (64). If the data met the assumption of normality, we conducted a one-way ANOVA to test for overall group differences, followed by Tukey’s *post hoc* test to determine which group means differed significantly (65). For data that did not meet normality assumptions, we used the Kruskal-Wallis test instead (66), followed by Dunn’s *post hoc* test for pairwise comparisons (67). For comparisons between two groups, we used the *t*-test (68) when both groups passed the normality test (64), and the Mann-Whitney U test when either or both groups did not pass the normality test (69).

## 3. Results

In this work, we used primary embryonic cortical neurons transduced with actin-GFP to study how the dynamics of actin change during development and when neurites interact with nanotopographic surfaces. In particular, we sought to examine actin dynamics on a timescale of seconds. Because it is known that the dendritic arbor of neurons changes significantly during development (26), we observed primary neurons between DIV 2 and 23 to determine whether actin dynamics changes as significantly as the dendritic arbor does during development. For neurons cultured on PDL-coated glass (control), we grouped our findings into early (DIV 2-5), middle (DIV 7-12), and late (DIV 14-23) stages. For neurons cultured on nanotopographic surfaces with 3.2 μm ridges, we grouped our findings into early (DIV 2-4) and middle (DIV 7-8) stages.

### Actin dynamics is persistent throughout different stages of development on both flat and nanoridged surfaces

To understand how actin-based motility varies during neuronal development and in response to substrate nanotopography, we first visualized actin activity under multiple conditions. Specifically, we sought to determine whether actin dynamics remains localized and transient, or shifts toward more directed, long-range behavior as neurons mature or encounter structural guidance cues. Figure 1 illustrates the cluster-tracking workflow’s ability to extract actin-wave dynamics from live-cell imaging videos for a range of developmental stages and substrate types. A representative mean-image from a neuron cultured on a flat surface (Fig. 1a-b) highlights an actin-flow event, which appears as a moving bright spot (arrows). The corresponding cluster tracks extracted from this video reveal localized movements with no apparent long-range flow or strong directional bias.

**Figure 1.**
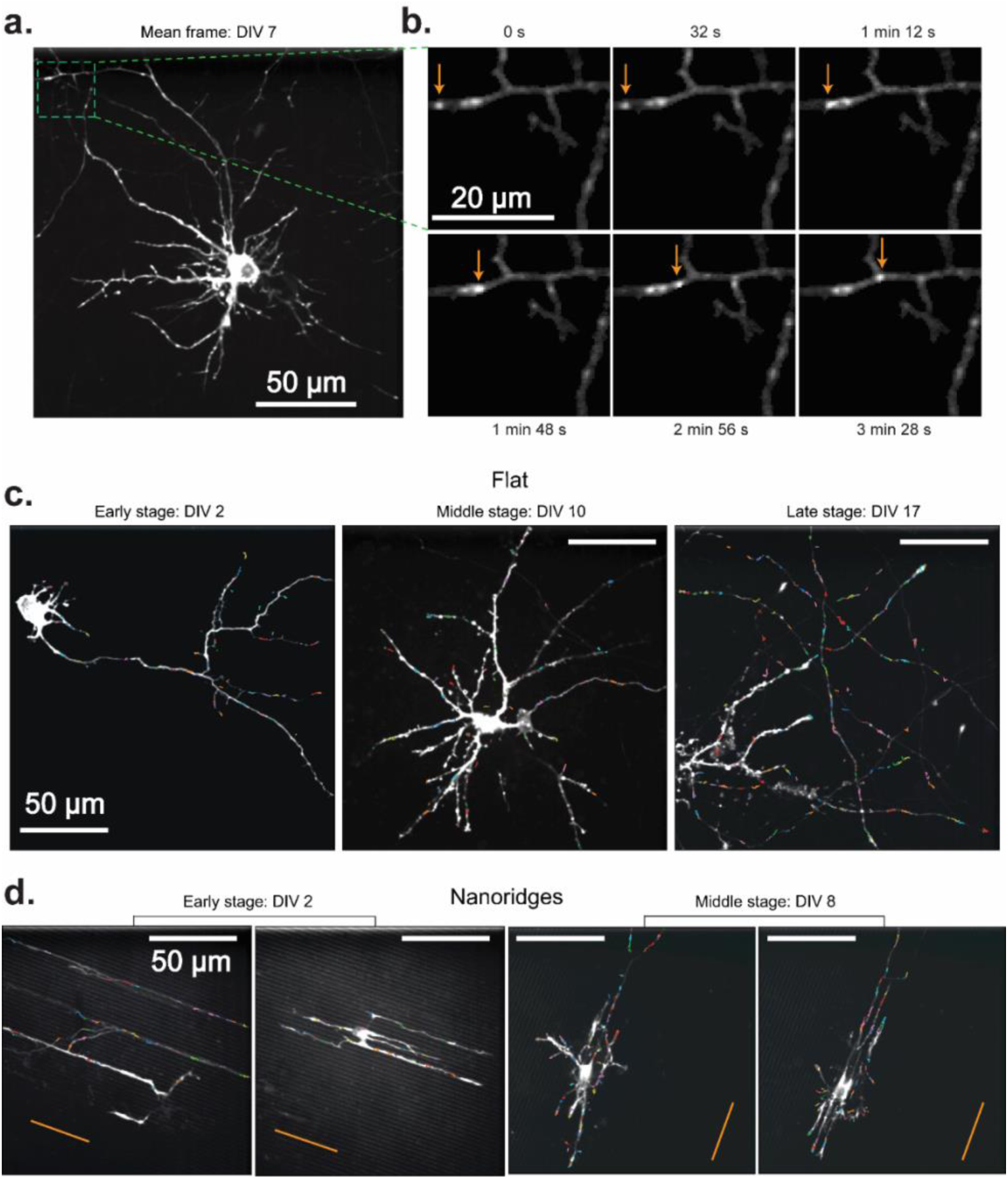
Actin waves in primary rat cortical neurons in different DIV stages on flat and nanoridged surfaces. (a) Mean image of a representative DIV 7 neuron cultured on a flat surface. The green box indicates a region of interest, which is shown in a magnified version in (b). (b) The trajectory of an actin wave (arrows) moving along a neuronal process. Actin tracks overlaid from representative videos, color-coded by individual track, for each developmental stage on (c) flat surfaces and (d) 3.2-μm-spaced nanoridges. On flat surfaces, the imaging durations were 5 min, 4 min 10 s and 8 min, respectively, for the representative videos from early, middle and late developmental stages. On nanoridges, the imaging durations were 5 min, 5 min, 10 min and 10 min, respectively, for the representative videos from early, early, middle and middle developmental stages. The orange lines indicate the nanoridge alignment.

These spatially confined actin events are observed consistently across developmental stages on flat substrates (Fig. 1c), and in early- and mid-stage neurons on nanoridged substrates (Fig. 1d), suggesting that the fundamental nature of actin activity remains transient (Suppl. Fig. 2) and spatially restricted (Suppl. Fig. 3) despite changes in growth conditions. Interestingly, in some cases, the cluster tracks appear to be aligned visually with underlying neuronal extensions, suggesting, qualitatively, that local cytoskeletal architecture influences the orientation of actin flow. Overall, the persistence of these short-range, dynamic actin events at different developmental stages and on different topographies (Suppl. Mov. 1-2) highlights the robust and adaptable nature of actin polymerization during neuronal growth and remodeling.

### Localized actin activity is less recurrent and less frequent on nanoridged substrates

Building on our observation that actin-polymerization events are brief and spatially localized at different developmental stages and on different substrates (Fig. 1), we next examined whether these events tend to recur in the same locations. Recurrence of activity in specific regions reflects spatial memory in the cytoskeleton, and suggests that certain sites undergo repeated remodeling. This spatial persistence is relevant for reinforcing cellular structures, guiding polarity, and directing neurite outgrowth. To test whether these properties are influenced by the cellular environment, we compared the recurrence and frequency of actin activity on flat and nanoridged substrates.

Figure 2 quantifies the recurrence and spatial distribution of localized actin tracks within neuronal processes. We first extracted representative optical-flow vectors from a single frame (Fig. 2a), and the clustered these vectors in space and time to generate robust actin tracks (Fig. 2b). We define mean recurrence as the fraction of actin tracks originating in the same spatial region in successive time windows, providing a measure of how consistently actin activity reappears in specific locations over time. Figures 2c and 2d explore how the mean recurrence depends on the spatial search radius and the temporal window size, respectively. We found that cells cultured on nanoridges consistently exhibited lower recurrence values compared to those on flat substrates (Fig. 2e), suggesting that substrate nanotopography disrupts the spatial persistence of actin waves. This trend was statistically significant at both the early (*p* = 0.00222) and the middle (*p* = 2.7 × 10^-5^) developmental stages, as assessed by Mann–Whitney U tests. Developmental stage alone, in contrast, did not significantly affect recurrence within the same substrate type (flat: *p* = 0.70; nanoridges: *p* = 0.271).

**Figure 2.**
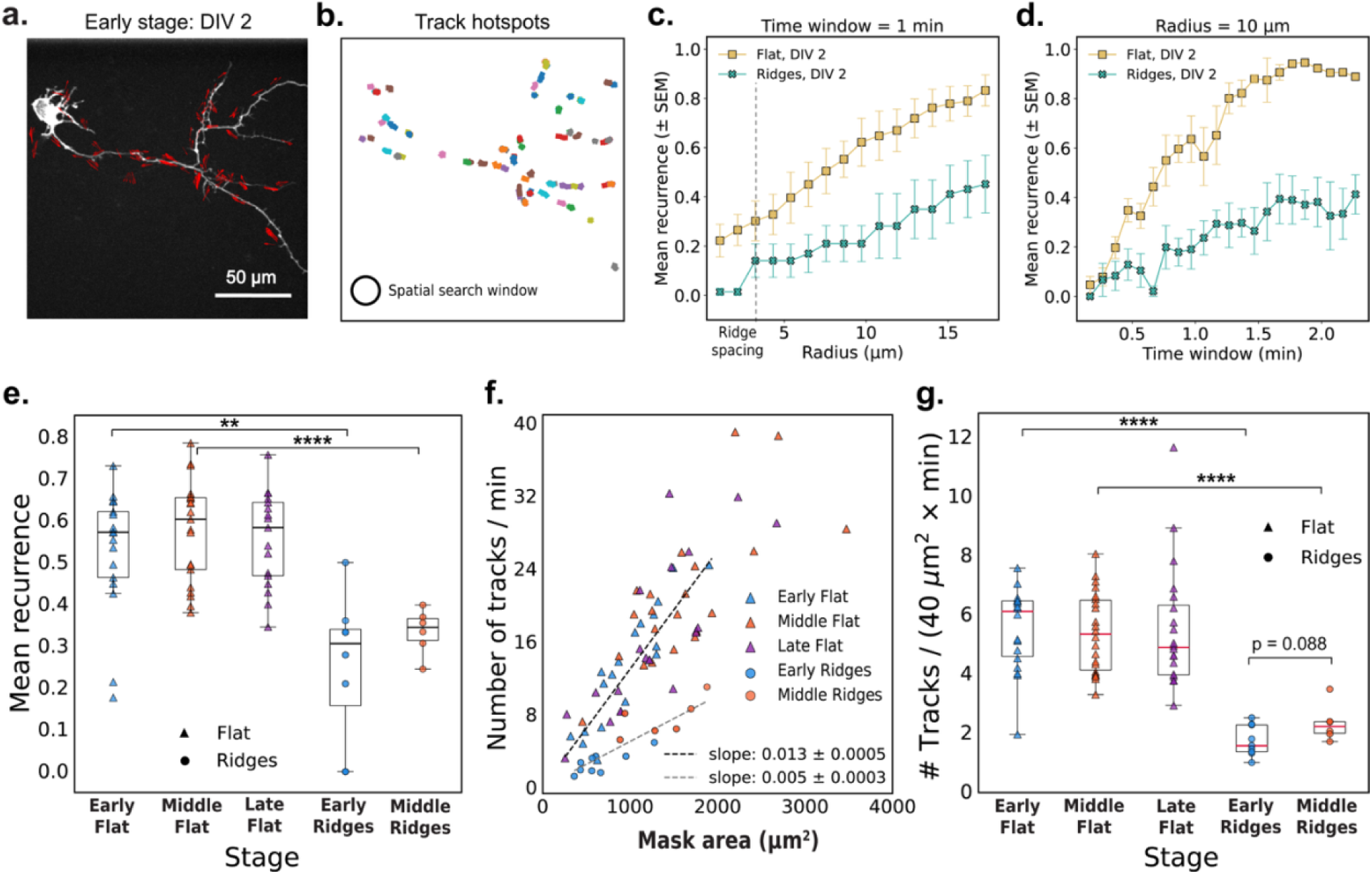
Actin-track dynamics and recurrence depend on the microenvironment. (a) Representative image frame with optical-flow vectors for a DIV2 neuron cultured on a flat surface. (b) Actin tracks observed over a 5-min interval from the same video as in (a). The circle illustrates the 10 μm search radius used to compute mean recurrence values shown in panels (d) and (e). (c-d) Mean recurrence quantifies the fraction of new actin tracks originating within a circular region during each time window. Mean recurrence as a function of (c) radius (with a 1-min time window) and (d) time window (at a fixed search radius of 10 μm) for two representative early-stage videos on flat and nanoridged substrates. The error bars represent SEM around the mean. (e) The mean recurrence for early (*N* = 17), middle (*N* = 21), and late (*N* = 17) videos, respectively, on a flat substrate and for early (*N* = 9) and middle (*N* = 6) videos, respectively, on nanoridged substrates. Search radius = 10 μm and time-window = 1-min (f) Tracks/min plotted as a function of the neuronal mask area (excluding the cell body) with linear fits (through origin) for flat (*N* = 55) and nanoridged substrates (*N* = 15). (g) Normalized track counts at different developmental stages. For (e)-(g), each scatter point corresponds to a unique video. Statistical tests were performed on pooled data. Significance is reported as *p*-value for 0.1 > *p* > 0.05, * for *p* ≤ 0.05, ** for *p* ≤ 0.01, *** for *p* ≤ 0.001, and **** for *p* ≤ 0.0001.

We next quantified the overall frequency of actin waves by measuring the number of tracks per minute and normalizing by the neuronal mask area. We observed a linear relationship between the total number of actin tracks per minute and the area of the neuronal mask for all conditions tested (Fig. 2f). This relationship is expected, as larger neurons have a broader cytoskeletal landscape, increasing the likelihood of detecting dynamic actin events. The linearity suggests that actin activity scales proportionally with cell area size, rather than being concentrated in a specific sub-region or regulated independently of cell area. Notably, the dependence of the frequency of actin waves on cell area was weaker in cells on nanoridges (slope = 0.005) compared to those on flat substrates (slope = 0.013). This trend is further supported by normalized track counts (Fig. 2g), which were significantly lower on nanoridges at both the early (*p* = 6.7 × 10^-5^) and middle (*p* = 1.1 × 10^-5^) stages. In contrast, no significant differences (flat: *p* = 0.67; nanoridges: *p* = 0.088), were observed across developmental stages for either substrate, suggesting that topographical surface exerts a stronger influence on actin dynamics than age.

Taken together, these results demonstrate that neurons not only initiate actin waves less frequently on nanoridges than on flat surfaces, but also exhibit reduced spatial recurrence of actin activity. These findings highlight the influence of physical nanotopography on the dynamics and spatial memory of cytoskeletal remodeling.

### Actin-wave-speeds depend on developmental stage and substrate environment

Next, we examined whether substrate topography also affects the speed at which these actin waves propagate. Actin-wave speed provides insight into how quickly polymerization-driven movements propagate through the cell, and reflects the timescales of cytoskeletal reorganization, mechanical resistance, and traveling signaling cues. Figure 3 illustrates how actin-wave speed varies with developmental stage and substrate. A representative time-lapse image from a DIV8 neuron on nanoridges (Fig. 3a) highlights a dynamic growth cone in which transient increases in fluorescence intensity mark actin activity over a 10-min period. Figure 3b shows alternating retraction and extension of this growth cone over time. This pulsatile behavior is typical of actin waves involved in guided protrusions *in vivo*. (8, 9, 23, 24).

**Figure 3.**
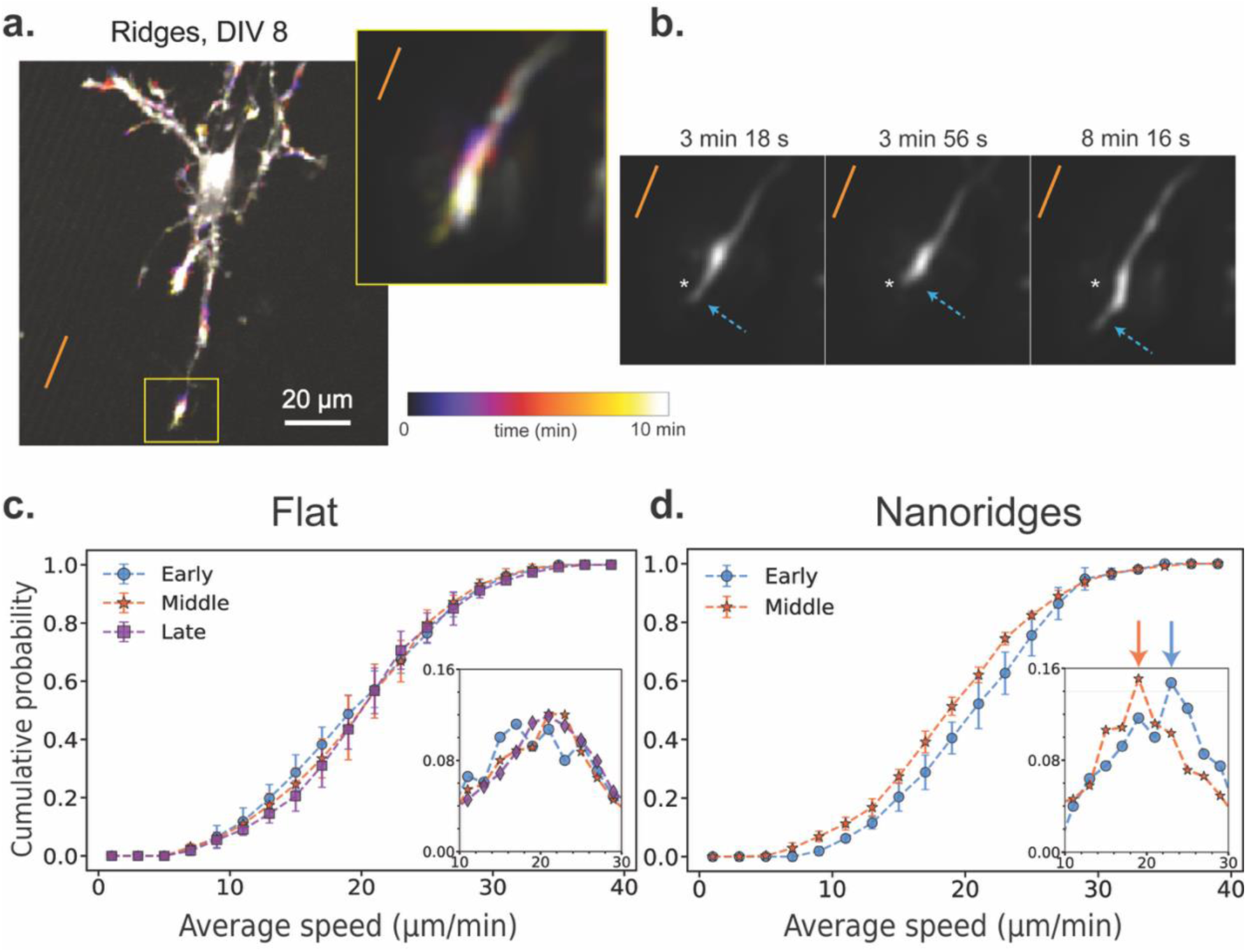
Actin-wave speed as a function of developmental stage and substrate identity. (a) A representative actin time-lapse image of a DIV8 neuron on a nanoridged surface. The inset shows a highly active growth cone over a 10-min period. The orange line indicates the nanoridge alignment. (b) The growth-cone trajectory from the same time-lapse image, showing alternating retraction and forward extension along the ridge direction. We applied the AI-based model ZS-DeconvNet (70) to denoise individual frames, exclusively for visualization purposes. The arrows show the locations of the leading edge of the growth cone. A fixed spatial reference point (asterisk) is shown in all three panels to aid visual comparison of the axon tip’s position over time. (c) CDF of the average wave speed for neurons cultured on flat surfaces for the three different developmental stages (*N* = 17, 21, 17 for the early, middle and late stages, respectively). (d) CDF of the average wave speed for neurons cultured on nanoridged surfaces for the early (*N* = 9) and middle (*N* = 6) developmental stages. In (c) and (d), the insets show the corresponding probability distribution function (PDF) near the peak. Number of bins = 20. The arrows in (d) point to the corresponding PDFs’ median values. The error bars represent the interquartile range of the CDF at each bin for all videos for a given condition, and the markers indicate the corresponding median values. For each pair of conditions, we calculated *W₁* between the median CDFs, as well as the maximum and minimum distances among all combinations of 25th and 75th percentile CDFs. These values are reported in the format: median (minimum to maximum). On flat surfaces, the *W₁* values were 0.53 (0.43 to 2.56) for early vs. middle, 0.54 (0.61 to 2.84) for middle vs. late, and 0.82 (0.66 to 2.91) for early vs. late stages. On nanoridges, the *W₁* value between early and middle-stage neurons was 1.59 (0.76 to 3.36).

To generalize these observations, we measured instantaneous actin-wave speeds under different conditions. On flat substrates (Fig. 3c), the speed distributions were similar for the early, middle, and late developmental stages, with an average of approximately 21 μm/min. However, on nanoridged substrates (Fig. 3d), a ∼10% decrease in wave speed was observed when going from the early to the middle developmental stage. We used the Wasserstein distance (*W₁*) to compare the distribution of speeds between stages and conditions. We found that the distance between early and middle stages on nanoridges was greater than that observed for any developmental comparison on flat surfaces. These findings suggest that neuronal maturation slows actin-wave propagation on nanoridged surfaces, whereas wave speed remains relatively constant on flat substrates. This observation implies that the mechanical or structural constraints imposed by nanotopography interact with developmental changes in the cytoskeleton, modulating how rapidly actin-based protrusions can extend. Such modulation can affect how neurons explore their environment and can stabilize emerging processes during growth.

### Cellular processes align actin waves more effectively than do nanoridges

Wave speed characterizes how rapidly actin structures move, but does not convey whether the waves propagate in a consistent direction or respond to local structural cues. To understand how actin dynamics is organized spatially, we next examined how actin tracks align with extracellular topography and with internal neuronal architecture. This analysis helps address whether external topography or internal cellular architecture plays a greater role in steering actin waves during neuronal development.

Figures 4a and 4c show 10 min of accumulated actin tracks for representative early- and middle-stage neurons cultured on nanoridges. Many of the tracks appear by eye to be aligned with the nanoridge direction. This visual impression is reinforced by zoomed-in regions (Fig. 4b), which reveal that both the actin tracks and neuronal processes are generally aligned with the nanoridges. To assess whether all actin waves follow a clear path, Fig. 4d highlights four representative middle-stage tracks that display prominent back-and-forth motion. These examples emphasize that many tracks are confined and lack a persistent direction, reinforcing the need to disentangle the roles of internal and external guidance cues. Figure 4e quantifies the alignment of actin tracks with nanoridges in the same representative early-and middle-stage videos shown in Figures 4a and 4c. Although most tracks follow the ridge direction, alignment is reduced in the middle-stage example.

**Figure 4.**
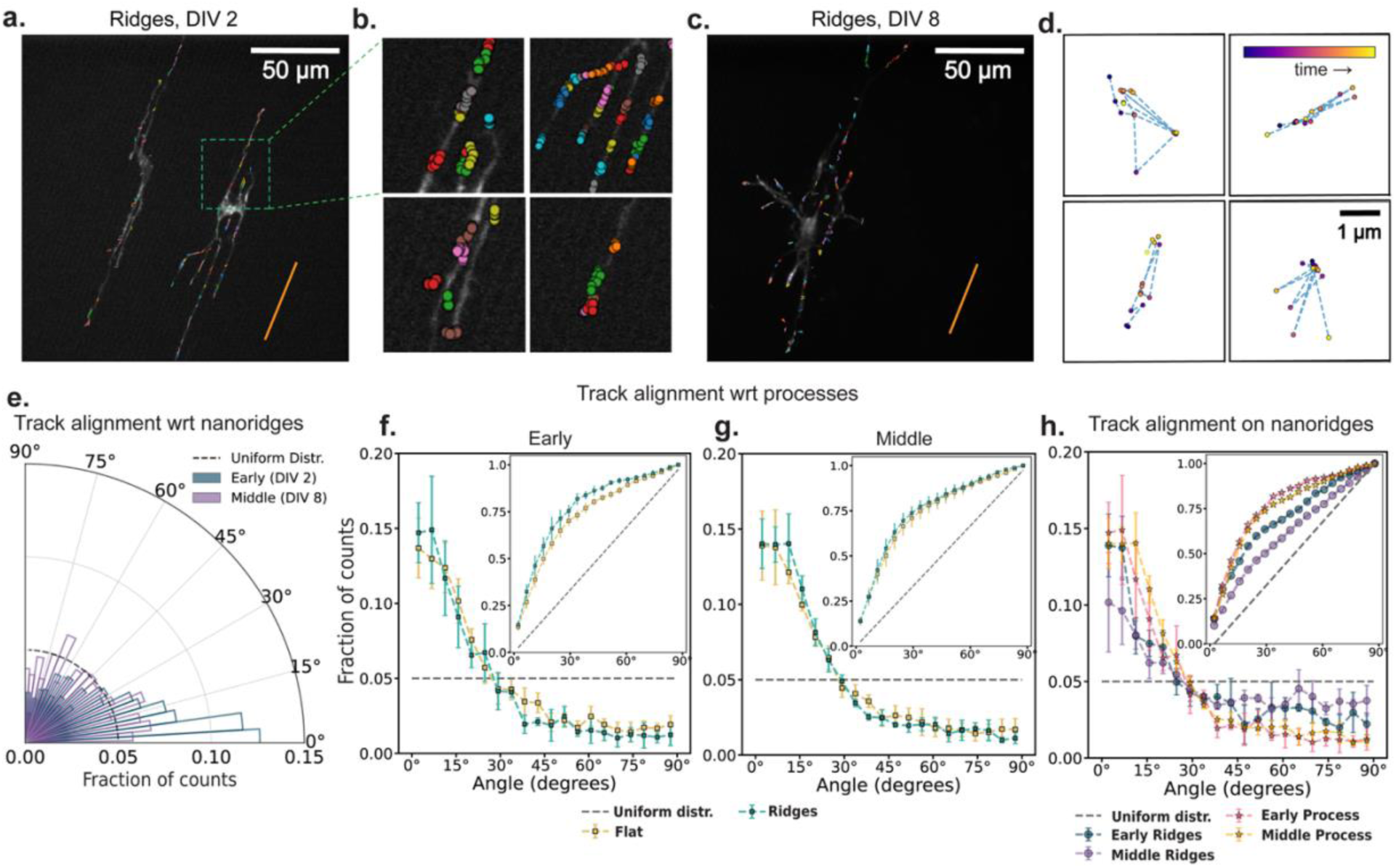
The influence of nanotopography and developmental stage on actin-track alignment. (a, c) Actin-cluster tracks accumulated for 10 min for two DIV2 and DIV 8 neurons respectively on nanoridges. The orange line indicates the nanoridge alignment. (b) Zoomed-in view of selected regions with actin tracks from (a). (d) Four representative actin tracks from (c) with frequent back-and-forth motion shown within fixed 4×4 µm windows. The data points are color-coded by time progression within each video (purple to yellow). (e) Alignment of actin tracks with underlying nanoridges in the same early (DIV 2) and mid-stage (DIV 8) videos, as shown in panels (a) and (c). The dashed line represents a uniform distribution of track alignments for the same bin size. There are 20 bins. (f, g) Fraction of track segments aligned at various angles with respect to neuronal processes on flat and nanoridged surfaces, at the (f) early and (g) middle developmental stages. (h) Relative alignment of track segments on nanoridges with respect to neuronal processes and the nanoridge for the early and middle developmental stages. The error bars represent the interquartile range of the PDF (or CDF in insets) at each bin across all videos for each condition, and the markers indicate the corresponding median values. From (f-h), the insets show the corresponding CDF. For each pair of conditions, we calculated *W₁* between the median CDFs and the maximum and minimum distances among all combinations of 25th and 75th percentile CDFs. These values are reported in the format: median (minimum to maximum). For early (f) and middle (g) stage neurons, the *W₁* value between flat and ridge surface movies were 3.91 (1.42 to 7.07) and 1.39 (1.34 to 7.03) respectively. On nanoridges, for early and middle stage neurons, *W₁* value between ridge and process guidance CDFs were 6.80 (3.0 to 13.5) and 12.07 (6.12 to 20.45) respectively. For process and ridge guidance comparisons, in early vs middle stage movies, the *W₁* values were 2.12 (1.16 to 5.96) and 7.23 (1.64 to 14.83), respectively.

We next explore the alignment of actin tracks with respect to the direction of the neural process in which they reside. At both the early (Fig. 4f) and middle (Fig. 4g) developmental stages, actin tracks remain strongly aligned with their respective processes, regardless of whether the neurons are cultured on flat or nanoridged substrates. A comparison of early and middle developmental stages on nanoridges (Fig. 4h) shows a clear reduction in track alignment with the nanoridges over time, while alignment with processes remains relatively stable. The inset in Fig. 4f further shows that track alignment with neuronal processes consistently exceeds alignment with nanoridges, suggesting that dynamic intracellular cues, such as membrane tension or cytoskeletal reorganization, may play a more dominant role in steering actin dynamics over time than does static external topography.

It is important to note that at early stages of development, the neuronal processes themselves are closely aligned with the nanoridges. This initial bias could give the impression that nanoridges directly guide actin dynamics, when in fact the processes may serve as the intermediate conduit. As development progresses, and the processes start diverging from the ridge direction, the decreasing alignment between actin tracks and nanoridges becomes more apparent. This observation raises a fundamental question: Is actin guidance influenced by the internal architecture of the cell versus the external physical environment, or does the physical environment guide the intracellular cues, and thus the actin dynamics of growing neurite tips, in a self-evolving feedback loop?

### Impact of nanoridges on growth-cone alignment

To explore the contributions of internal vs. external physical cues to cytoskeletal guidance, we next consider the structural features of developing growth cones. Specifically, we analyzed how the growth-cone alignment is related to nanoridge orientation over the course of development. Figures 5a and 5b show representative examples of growth cones that are aligned either parallel (at DIV 2) or nearly perpendicular (at DIV 8) to the nanoridges. The growth cone in Fig. 5b, marked by the green box, is especially notable for originating from a neighboring neuron located outside the field of view, and extending along a neurite that crosses multiple nanoridges. Thus, by later stages of development, neurites are capable of growing across the ridges, potentially enabling more diverse growth-cone orientations. In Figure 5c, we show the growth-cone alignment as a function of growth-cone width, revealing a developmental shift. At early time points, growth cones display a strong preference for alignment along the nanoridges. This preference decreases as development progresses.

**Figure 5.**
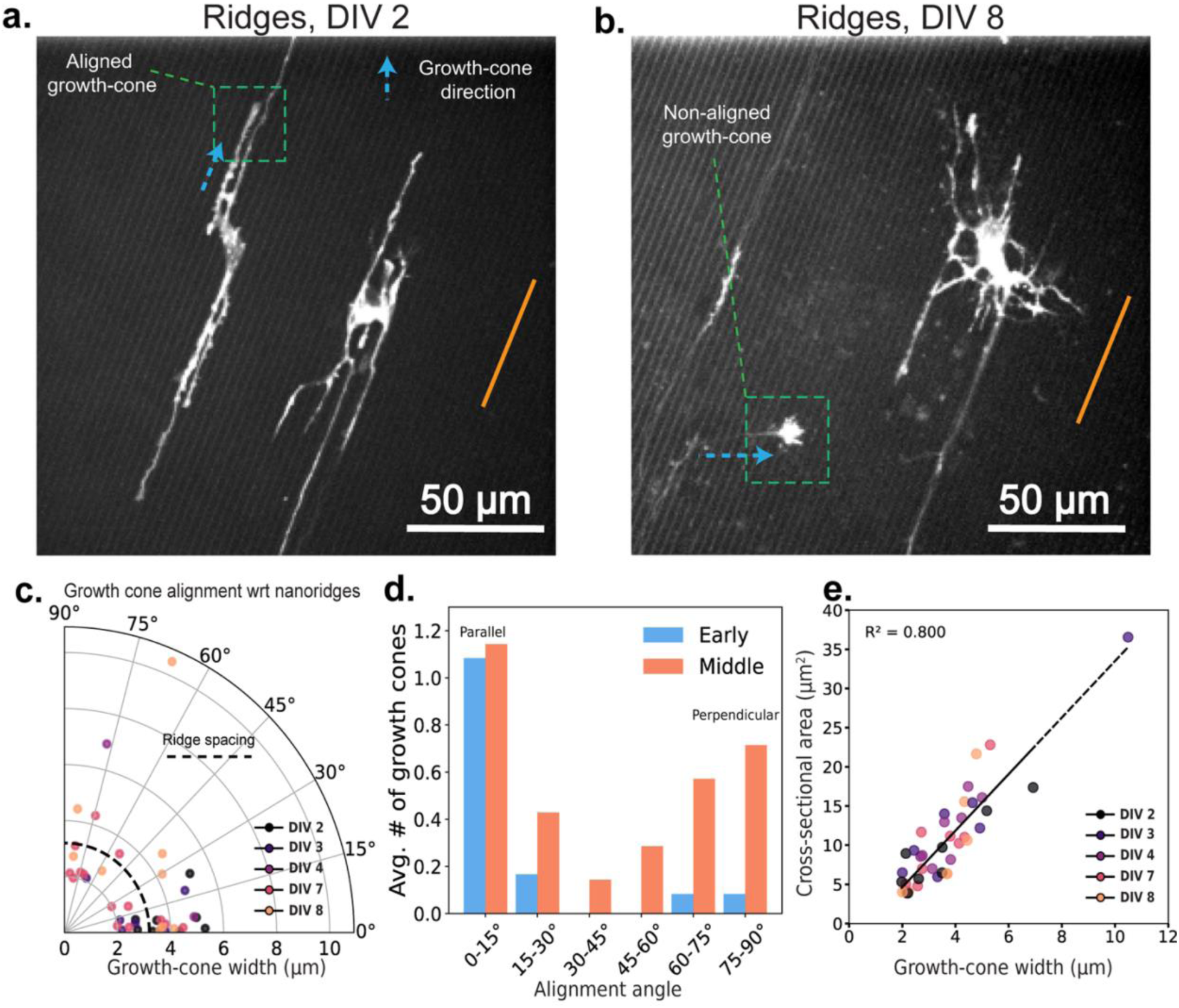
Growth-cone orientations at different developmental stages for neurons on nanoridges. Representative growth cones oriented (a) parallel to (DIV 2) and (b) perpendicular to (DIV 8) the nanoridges. The nanoridge directions are indicated by orange lines, and the growth-cone orientations by arrows. (c) The distribution of growth-cone alignment angles with respect to the nanoridge as a function of the growth-cone width. Darker to brighter colors indicate neuronal development stages from DIV 2 to DIV 8. (d) The average number of growth cones per neuron as a function of the alignment angle with respect to the nanoridges for early-and middle-stage neurons. Growth cones at angles between 0°and 15° are considered to be parallel, whereas growth cones at angles between 75*°* and 90*°* are considered to be perpendicular. (e) The relationship between the cross-sectional area and width of the growth cones, color-coded for developmental stage. Videos with at least one growth cone were considered for analysis. *N* = 7, 4, 1, 4, 3 for DIV 2, 3, 4, 7, and 8, respectively. Due to low *N*, we did not run statistical tests on the average number of growth cones across DIV stages and alignment angles.

We also observe that growth cones rarely adopt intermediate orientations around 45°, instead favoring directions that are either approximately parallel to or nearly perpendicular to the nanoridges (Fig. 5d). Although the average number of growth cones aligned parallel to the ridges remains relatively consistent over time, there is a substantial increase in the number of perpendicularly aligned growth cones during later stages.

This bimodal distribution suggests a shift in the balance of guidance mechanisms over time. Early in development, growth cones appear to align closely with the nanoridges, indicating that substrate nanotopography serves as a primary external cue. As neurons mature and more neurites populate the surface, spanning across multiple ridges, the physical landscape becomes increasingly complex. In this setting, new growth cones may interact not just with the substrate, but also with these pre-existing neurites, which create internal mechanical and geometric constraints.

Interestingly, we also observed perpendicular growth in areas lacking visible pre-existing neurites, implying that additional factors, such as developmental changes in adhesion or cytoskeletal dynamic, may be at play. Similar dual-mode guidance patterns have been reported previously (47, 71, 72). These shifts likely lead to a transition from continuous to more discrete preferred growth orientations, contributing to the observed bimodality. One possible mechanism is that new growth cones follow earlier neurites that have already crossed the ridges. These existing neurites may exert mechanical tension while straightening during development, promoting a greater amount of perpendicular orientation. Alternatively, geometric constraints, such as filopodia reaching the base of adjacent ridges, may pull the growth cone downward, reinforcing perpendicular growth.

Finally, Figure 5e shows that although growth-cone cross-sectional area is positively correlated with width, neither area nor width changes significantly with developmental stage. This observation indicates that size alone does not determine alignment or guidance behavior. Instead, it is the evolving balance between external substrate features and internal structural constraints that shapes how growth cones navigate their surroundings over time.

## Discussion

We have characterized actin dynamics in primary rat cortical neurons at developmental stages ranging from DIV 2 (early) to DIV 23 (late), and we have quantified these dynamics using an optical-flow based technique. Because primary cells have a finite lifetime *in vitro*, we focused on time points at which our primary cultures were healthy (assessed twice weekly during media changes) and used for experiments when successful transduction was possible. In particular, because our nanotopographic surfaces were not optimized for longer-term (DIV 10+) neuron survival, we only studied neurons on ridges during the early and middle developmental stages.

An additional consideration for the interpretation of our findings is the presence of glial cells in primary neuronal cultures. Recent studies have shown that glial cells can affect neurite guidance (73), and likely also actin dynamics. We did not control for the chemical or growth factors released by either neurons or glial cells, both of which can affect neurite guidance (74). Moreover, the presence of glial cells modulates the physical environment (75), which may affect neurite guidance and actin dynamics. Future work will dissect the complex interplay between the mechanical and chemical cues that drive actin dynamics and the consequences of eliminating glial cells from the *in vitro* environment.

Because actin dynamics can serve as a primary sensor of physical signals (46, 52, 53), we assessed how a well-established physical guidance cue, nanoridges, affects actin dynamics and neurite tips. Most previous studies (8, 23, 24, 76–78) used imaging frame rates on timescales of minutes, which is too slow to capture the actin-polymerization waves that act as sensors of the physical environment. By imaging the dynamics of actin at a rate of 0.5 frames/s, we were able to analyze how polymerization and depolymerization waves in primary neurons contribute to growth-cone development, neuronal maturation, and sensing of the physical microenvironment.

We demonstrated that, on flat surfaces, the actin-wave speeds remain constant throughout neuronal development. In contrast, on nanoridges, actin-waves are suppressed, and the wave speed decreases slightly with increasing neuronal maturity. These findings suggest that actin is indeed functioning as a sensor of the environment, as we found in our previous work on astrocytes (53): when the environment is not as physiologically relevant (flat, PDL-coated glass), actin wave speed does not change as neurons mature, but when the environment is more realistic (nanoridged substrate), the actin-wave speed decreases with neuronal maturity. It is possible that the differential effects on speed observed on flat vs. nanoridged substrates are related to the more physiologically-relevant environment of the nanoridged substrates. Moreover, neurite orientation appears to be the primary guidance cue for the direction of actin-wave propagation throughout development. As neurons mature, the guidance of actin by nanoridges is reduced, and both neurite-tip and growth-cone guidance by nanoridges diminish, with an increase in motion perpendicular to the nanoridges for actin waves, neurite tips, and growth cones.

It would be interesting in future work to explore, via pharmacological experiments, whether branched or linear actin contributes more to the guidance we observe. Previous studies have generally indicated that branched actin is responsible for swellings in dendrites, whereas linear actin is responsible for filopodia extension (79). A delicate balance of both branched and linear actin is necessary for axonal patterning (80, 81). Indeed, a recent preprint studying contact guidance in epithelial cells suggests that formins (regulators of linear actin) increase contact guidance, whereas Arp2/3 (a regulator of branched actin) decreases contact guidance (82). In our system, it is possible that in younger neurons there is more formin activity (and therefore more guidance by nanoridges) and that in older neurons there is more Arp2/3 activity (and therefore less guidance by nanoridges). Possibilities for future work include dissecting the roles of branched vs. linear in nanoridge guidance and exploring the spatiotemporal relationships between actin dynamics and dynamics of the signal-transduction network in primary neurons (13, 15, 16, 18–20).

Our findings demonstrate that the ability of neurons to sense and follow physical guidance cues may change with developmental stage, and raise the question of how such a change in physical sensing capabilities could impact brain development and regeneration. Recent studies have shown that promoting actin dynamics as a sensor of the physical microenvironment can enhance axon regeneration in the adult nervous system, particularly through the regulation of specific actin-binding proteins such as ADF/Cofilin (7). Understanding how nanotopography influences actin dynamics during various stages of development may thus provide valuable insights into potential strategies for promoting axon regeneration after injury or disease. It is important to note that our conclusions are limited to those that can be derived from an embryonic system, because even the “late stage” neurons in our study are still young compared to the adult neurons in a living organism.

Finally, our findings may have implications for the development of novel therapeutic approaches for processes that are known to involve actin dynamics, in particular neuroregeneration (83) and the treatment of neurodegenerative diseases (84). Many studies have shown that brain stiffness decreases with age (83–86), and that varying substrate stiffness affects neuronal morphology (36, 37). Thus our work provides a baseline for future studies investigating how to tune actin dynamics and thus neurite tip guidance to physical microenvironments that more closely mimic the brain at various developmental or disease stages.

## Acknowledgments

This work was supported by the US Air Force Office of Scientific Research MURI grant FA9550-16-1-0052 to W.L. and J.F. It was also supported in part by funds from the Intramural Research Program of NINDS, NIH: ZIA NS003013 to E.G.

